# Homeorhetic regulation of cellular phenotype

**DOI:** 10.1101/2025.06.06.658216

**Authors:** Manish Yadav, Daniel Koch, Aneta Koseska

## Abstract

How cells translate growth factor (GF) signals into context-specific phenotypes remains a fundamental question in cell biology. The classical view holds that cells translate a constant concentration of a GF with specific chemical identity to a steady state activation of the underlying signaling pathway, resulting in a defined phenotypic response. However, recent findings suggest that even a single GF, when presented in a pulsatile manner, drives differential phenotypic responses depending on the frequency of stimulation. To reconcile these views, we introduce a novel conceptual framework of “*signaling homeorhesis*”. Unlike homeostasis, homeorhesis describes the stable evolution of signaling trajectories over time. Defining this concept quantitatively using a dynamical systems framework, we use mathematical models of the Epidermal Factor Growth Receptor (EGFR) and the Tropomyosin receptor kinase A (TrkA) networks, as well as available experimental temporal protein activity recordings in PC-12 cells to demonstrate that cells classify GF signals to unique signaling trajectories that encode for distinct cell phenotypes, irrespective of the GF identity. We thereby propose that the cellular phenotype is determined in real-time, as the cell actively interprets the growth factor signals from its environment.

## Introduction

Signaling networks in the individual cells of a developing organism or an adult tissue continuously sense and process chemical signals secreted by the neighboring cells in their environment and transduce it to immediate early gene (IEG) networks, ultimately forming corresponding phenotypic responses^1–8^. Since these chemical signals are noisy and vary in space and time, the signaling and the IEG networks likely integrate the information obtained over time, suggesting that both the encoding and decoding of the information must be robustly and reproducibly performed *in real time*, before the networks approach a steady state. Considering the mapping of a chemical signal to a phenotypic response as a process of computation performed by the biochemical network, an accurate interpretation of the signals relies on *signaling specificity*, as one ligand can generate differential cellular responses even by activating the same downstream components^9,10^. At the same time, the network should enable an appropriate *signal classification* such that different chemical inputs can induce the same phenotypic response^9,11^. Therefore, understanding how this mapping is performed by biochemical networks in single cells is a fundamental problem in signal transduction.

Among the most prominent and studied examples of how signaling specificity is efficiently achieved by activating shared components when cells are stimulated with constant signals, are the responses of the ERK family of MAP kinases to different growth factors^6,9,11–14^ and the cAMP signaling in cardiac myocytes^15–17^. It has been suggested that possible mechanisms include protein scaffolding^18–22^ and nano-domain signaling^23,24^, as well as pathway-specific sensing of different ligand features such as their molecular identity and concentration^14,25^. In the majority of these cases, the biochemical network is assumed to have a modular, pathway-like structure, and specificity is thought to arise through the underlying feedbacks, or by cross-inhibition between pathways, leading to the stabilization of a specific signaling steady state that the gene regulatory networks (GRNs)presumably decode to a specific cell fate. However, it has been experimentally shown that phenotypic responses cannot be effectively predicted from the steady-state level of proteins within a single pathway^10^. This, in turn, suggests that the encoding of the information is distributed throughout the large-scale protein-interaction network, rather than the network being modular, which is in line with proteomics studies demonstrating a complex, highly interconnected intracellular protein-interaction network^26–28^.

In a pioneering study, stimulating cells with a pulsatile GF in microfluidic devices, Ryu et al.^11^ have demonstrated experimentally that a single growth factor can bias towards a specific cell phenotype, i.e., proliferation or differentiation, only by changing the stimulation frequency. In particular, the authors have shown that stimulating rat adrenal phaeochromocytoma cells (PC-12), a classical model system to address how specific functionality arises from the operation of the MAPK signaling cascade^9,12,19^, with Epidermal growth factor (EGF) with 3^′^3^′^ (3min pulse, 3min inter-pulse interval) or 3^′^60^′^ induced proliferation, whereas stimulating with 3^′^10 and 3^′^20^′^ induced PC12 cell-differentiation. Similarly, stimulation with Nerve growth factor (NGF) with 3^′^3^′^, 3^′^10^′^ and 3^′^20^′^ induced differentiation, whereas only stimulation with a 3^′^60^′^ signal profile induced proliferation. Importantly, it was demonstrated that cells perform an accurate interpretation of the time-varying growth factor signals, despite the high degree of cell-to-cell variability in the single-cell protein activity profiles^11^. Very recently, a similar observation has been made for HEK293T cells and H9 human embryonic stem cells^29^. These results thus suggest that cells do not only rely on the molecular identity of the signal to guide their phenotypic responses but rather interpret the temporal distribution of the signals they receive over time. This, however, challenges traditional signaling concepts and reopens the question of how the cellular phenotype is regulated and determined, especially when the growth factor signals are not constant over time. Rephrasing this from a computational perspective, one can ask how distributed biochemical networks generate *specificity* in signaling such that dynamic growth factor signals can be mapped to distinct phenotypic responses, while at the same time enabling *signal classification* in the sense that chemically distinct growth factor signals can generate equivalent phenotypic responses. Therefore, the necessity arises for a novel conceptual framework that can explain how phenotypic responses are regulated by cellular signaling.

Here, we introduce a quantitative concept rigorously based on the underlying non-linear and non-autonomous dynamics of signaling networks to describe signaling regulation during phenotypic determination in the presence of dynamic signals. We tackle this question from the aspect of cellular computations by bringing formally together three abstractions that exemplify a general approach to the problem. First, when facing dynamic growth factor signals, the cell phenotype is determined by the signaling trajectory or the change of the state of the distributed signaling network in time; second, time-varying signals encoding for a specific phenotype, irrespective of the chemical identity, map to a well-defined family of signaling trajectories, and signaling trajectories encoding for different phenotypes diverge over time. Third, the cellular phenotype is implicitly encoded in the temporal evolution of the trajectory in signaling space, and it can be directly read out and decoded in real-time by the IEGs, much earlier than the network sets to a steady state.

We hereby propose that signaling *homeorhesis* determines cell phenotype in dynamic environments. In contrast to the homeostatic view^30,31^, by which the system maintains balanced states and returns to a static steady state after intermittent perturbations, the homeorhetic view describes the maintenance of a dynamic trajectory, which, as we show here, is uniquely determined by the time-varying external signals in conjunction with the molecular system on which they act. Following the description of the formal concepts and its quantitative implementation for a generic signaling networks, we illustrate and validate our framework based on experimental time series data from PC-12 cells along with simulations of the Epidermal Factor Growth Receptor (EGFR) and the Tropomyosin receptor kinase A (TrkA) networks, showing how cell fate specification in PC12 cells arises via homeorhetic signaling regulation.

### Mathematical framework of signaling trajectories

We introduce a mathematical framework for the description of signaling trajectories, consistent with the existence of a distributed, highly-interconnected intracellular protein-interaction network^26,27^. Theoretically, when the inputs are temporally complex, the protein-interaction network operates as a complex system whose dynamics is driven by the external inputs that constantly impinge on it. This means that the dynamics of the signaling network cannot be separated from the environment in which it operates and is, therefore, determined by the underlying network topology in conjunction with the external signals (Figure 1a). The evolution of the overall signaling state of the network is thus given by:

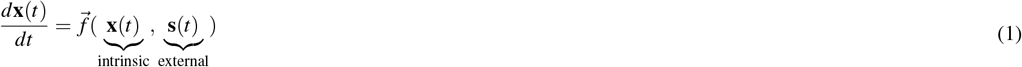

where the scalar *t* ∈ ℝ = (−∞, +∞), **x** ∈ ℝ^*n*^ is the internal state of the system, **s** is the vector of the exogenous signal. As the dynamics of the system explicitly depend on the temporal evolution of the external signal, this framework is formally referred to as *non-autonomous system’s description*^32^. To keep molecular details minimal and focus on the general dynamic principles, we adopt a canonical modeling framework from biochemical systems theory^33,34^ such that for a biochemical network with *i* = 1, ..*N* components, Eq. 1 takes the general form:

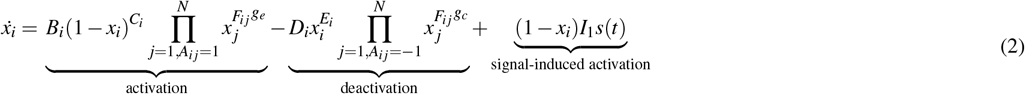

where *B,C* / *D, E* represent the autonomous activation/deactivation rate constants, *I* - signal-induced activation, and 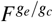 - the rate of activation/deactivation of *i*^*th*^ by the *j*^*th*^ component, if there exists a link between them i.e *A*_*i j*_ ≠ 0, whereas the *A*_*i j*_ ∈ [−1, 0, 1] is the connectivity matrix of the arbitrary signaling network. To identify the solutions of the non-autonomous system (Eq. 2), it is necessary to consider a family of solution operators 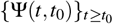^32, 35^ that depend both on the final and the initial state. These solutions are, therefore, complete trajectories. By definition, the map *x* : ℝ → ℝ^*m*^ is a complete trajectory if

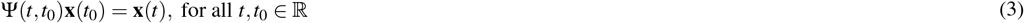

**Figure 1.**
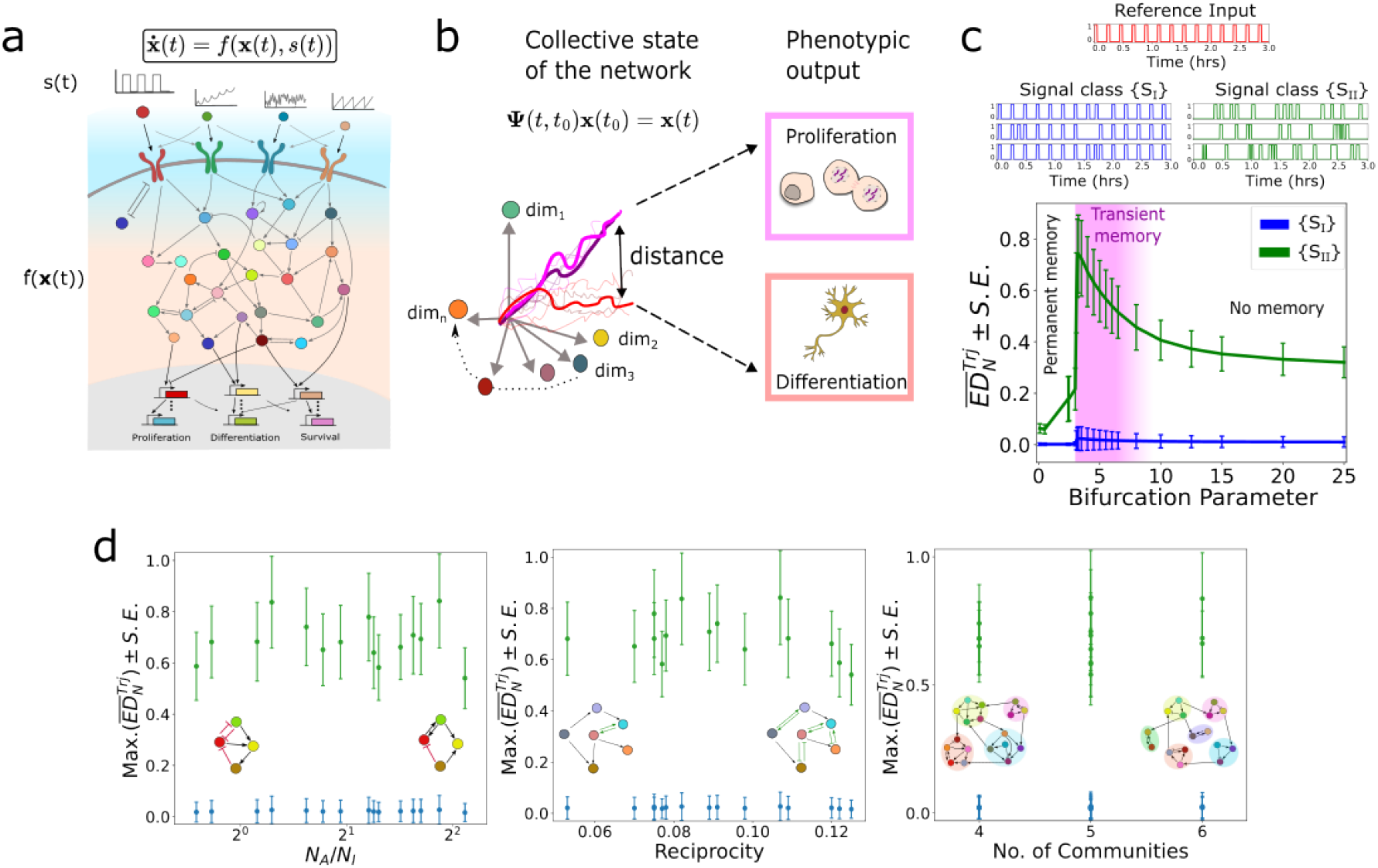
Basic features of homeorhetic regulation of signaling. (a) Schematic diagram of a distributed biochemical network with an arbitrary topology processing external dynamic stimuli. *s*(*t*) denotes the signal profile, and *f* (*x*(*t*)) - the signaling network dynamics. (b) Schematic representation of families of signaling trajectories (Ψ), depicting the evolution of the overall signaling state of the network over time. Families of signaling trajectories encoding for different phenotypes diverge in the signaling state-space, as denoted by an increased distance. (c) Classification of signals based on their temporal profile by a distributed biochemical network (*N* = 30) operating at three different dynamical regimes. Input signal classes with different frequency profiles are generated from the reference input (red, 3^′^10^′^ - 3min input, 10min inter-pulse interval), such that {*S*_*I*_} (blue) and {*S*_*II*_} (green) correspond to shuffling the reference pulses with probabilities 0.95 and 0.1, respectively. Average 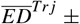 standard error between trajectories generated by 10 distinct inputs from {*S*_*I*_} and {*S*_*II*_} respectively, with respect to the *reference* trajectory in the 30-dimensional phase-space is shown as a function of the bifurcation parameter (receptor concentration). See also Supp. Figure 1. (d) Maximum value of 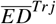 for organization at the critical transition between no memory and permanenet memory, estimated for (left) different ratios of activatory to inhibitory links (*N*_*A*_*/N*_*I*_), (middle) different network reciprocity values and (rigth) different network’s community numbers. The average ± standard error from 15 different network topologies, estimated for 10 signals from {*S*_*I*_} and {*S*_*II*_}, is shown.

Thus, in contrast to the classical autonomous systems description, where the signaling state of the network is determined only by the underlying network topology and the resulting steady states, in non-autonomous systems, the system’s behavior is defined by the trajectory, or the change of the state of the system (Figure 1b). For an *N*− dimensional signaling network, at each point in time, the signaling state is represented by an *N*−dimensional vector in the state-space - the space spanned by the system’s variables, which in this case are the concentration of the protein activities of the network. The signaling trajectory is thereby a set of vectors that describes the evolution of the state of the signaling network over time.

This substitution of the notion of steady-states for autonomous systems with the notion of complete trajectories for non-autonomous systems intuitively follows from the fact that when a system is subjected to time-varying signals, either the position, shape, and size of the steady-states change over time (state-space geometry) or their number and stability (state-space topology)^36–38^. In an abstract sense, the same notion has been schematically depicted by Waddington^39^, where the shape of the landscape (resembling the state-space) is continuously modulated by the pull of the strings (equivalent to time-varying external signals), which in turn determines the trajectory of the rolling marble which not necessarily needs to set to a defined valley (steady state).

### Rationale of homeorhetic regulation of the cellular phenotype

Having defined signaling trajectories, which are formal solutions of a biochemical network responding to time-varying growth factor signals, we introduce the concept of *homeorhesis*. First, we define the maintenance of a stable family of signaling trajectories that evolve over time as *signaling homeorhesis*. Based on this definition, we propose that each family of signaling trajectories distinctively defines a unique phenotype, a process which we term *homeorhetic regulation of cell phenotypes*. As each family is uniquely determined by the time-varying external signals in conjunction with the biochemical network on which it acts, this hypothesis has two major implications: *i)* families of signaling trajectories encoding for different phenotypic outputs diverge over time in the signaling state-space (Figure 1b), thereby ensuring specificity in signaling even if the network does not consist of modular pathways; and *ii)* growth factors with different chemical composition can be mapped to the same family of signaling trajectories, generating equivalent phenotypic responses.

To explore these concepts, we utilized a generic receptor model^38^ coupled to system Eq. 2 to study the response dynamics of a paradigmatic signaling network to time-varying input signals. We generated an *N* = 30 dimensional signaling network with arbitrary network topology and reaction parameters (cf. Methods). Following our recent findings that temporal memory in receptor networks aids the integration of spatial-temporal signals^38,40,41^, we hypothesized that such temporal memory may play an important role in signaling homeorhesis. We thus parameterized the core receptor circuit, and thereby the signaling network, to operate in three distinct dynamical regimes: no memory, permanent memory, and temporal memory, corresponding to the organization in a monostable regime, bistable regime, or at the critical transition between mono- and bistable regime, respectively. The organization of the network in these three distinct dynamical regimes relies on the receptor concentration on the membrane, which here serves as a bifurcation parameter (Figure 1c). To study if this minimal model of a signaling network gives rise to signaling homeorhesis, we constructed a *reference* temporal signal with an arbitrarily-chosen, fixed inter-pulse interval between any two consecutive pulses (Figure 1c). We then generated two signal classes, {*S*_*I*_}, {*S*_*II*_}, each containing *N*_*I,II*_ = 10 realizations, such that the Euclidean distance between the reference signal and each signal from the {*S*_*I*_} class was ∼ 0, and ∼ 1 to the signals from {*S*_*II*_}, implying different levels of similarity with respect to the reference signals. For each of the generated signals, the evolution of the network’s dynamics over time that describes a signaling trajectory was determined, and the Euclidean distance 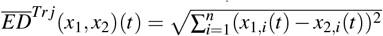, to the reference signaling trajectory was estimated. The 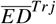 between the reference signaling trajectory and the signaling trajectories induced by signals from {*S*_*I*_} remained close to zero irrespective of the presence of memory in the system, implying that signals with a similar temporal structure are mapped to a unique family of signaling trajectories, i.e. the overall signaling activity patterns of the network across time remain almost identical. On the other hand, the signaling trajectories induced by signals from {*S*_*II*_} diverged in state space from the trajectory of the reference signals, with the separation being most pronounced (highest 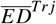) when the signaling network is characterized by temporal memory (Figure 1c). This suggests that signals with distinct temporal profiles are mapped to different families of signaling trajectories, i.e., the overall activity pattern of the signaling network over time is distinct, thereby achieving signaling specificity even when the same distributed network processes the signals. Thus, temporal memory corresponding to a prolonged phosphorylation of the receptor afetr signal removal^38^, is crucial for the distinct integration of the signals depending on their frequency profile.

Next, we sought to identify how the network properties influence signaling homeorhesis when the receptor module is characterized by temporal memory. As a first step, we drastically reduced the dimensionality of the signaling network, keeping only the core receptor module (*N* = 3). Interestingly, we found the classification of the signals into trajectory families to be preserved, however, the separation between the trajectory families was significantly decreased (Supp. Figure 1). We next tested the effect of the network topology (*N* = 30 node network, Figure 1c) by systematically varying *i)* the number of activatory and inhibitory links (*N*_*A*_*/N*_*I*_) in the network, *ii)* the network’s reciprocity, representing the likelihood that nodes are mutually linked, and *iii)* the number of communities, representing the subset of nodes among which the connections are denser than in the rest of the network. The distance between the trajectories of class *I* and the reference signal’s trajectory remained ∼ 0, in contrast to the 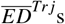 from {*S*_*II*_} signals that were maintained *>* 0.5 accross all network modifications (Figure 1d). These results suggest that signaling homeorhesis likely emerges due to the presence of temporal memory in the signaling network without the requirement for a specific network topology. However, high dimensionality of the signaling network is necessary for reliable and robust separation of the signaling trajectories, thereby leading to signaling specificity.

### Differential fate of PC12 cells arises through signaling homeorhesis

We next explored experimental evidence for homeorhetic regulation of cellular phenotypes, focusing on the PC12 cells paradigm described in the introduction. Experimental evidence shows that EGF stimulation with 3^′^3^′^ or 3^′^60^′^ induces proliferation, whereas stimulation with 3^′^10^′^ or 3^′^20^′^ leads to differentiation^11^. The EGF, on the other hand, activates the Epidermal Growth Factor Receptor network (Figure 2a), for which we have previously experimentally demonstrated the existence of transient memory with an average duration of ≈ 40min in single cells^40,43^. This experimental paradigm, therefore, represents an ideal example for testing homeorhetic regulation of cell phenotype, as it fulfills the necessary conditions outlined above.

**Figure 2.**
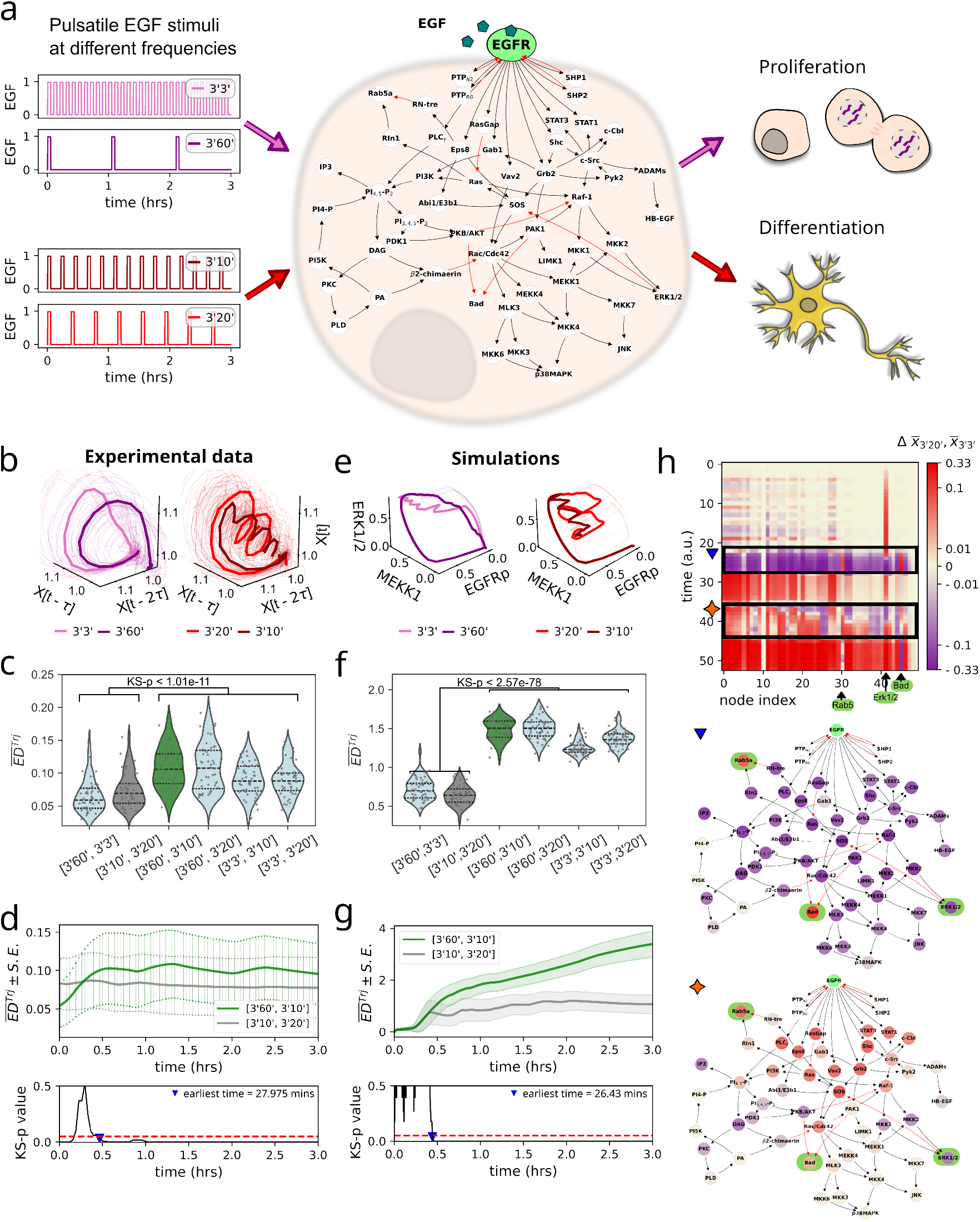
Homeorhetic regulation determines phenotypic responses of PC12 cells to time-varying growth factor signals. (a) Network structure of experimentally identified EGFR network^42,43^. Schematic depicting differential phenotypic response of PC12 cells, proliferation and differentiation, when stimulated with 3^′^3^′^ and 3^′^60^′^ or 3^′^10^′^ and 3^′^20^′^, respectively^11^. (b) Reconstructed signaling trajectories from experimentally obtained temporal ERK phosphorylation profiles^11^, depicting the overall evolution of the signaling network dynamics over time for different EGF input frequencies. Thin solid lines: signaling trajectories reconstructed from single cell profiles, thick solid lines: average signaling trajectories per input frequency. Note the similarity in the shape of the average signaling trajectories corresponding to a defined phenotype. (c) Euclidean distance 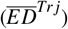 between signaling trajectories reconstructed from *N* = 30 cells per condition, calculated for the first hour of stimulus presentation, for all combinations of signals belonging to the same class i.e, proliferation and differentiation, and across the two classes. Trajectories were initially time-warped (Methods). (d) Evolution of the mean 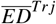 calculated between signaling trajectories reconstructed from *N* = 30 single cells corresponding to the same class (3^′^10^′^ and 3^′^20^′^, grey), or different class (3^′^60^′^ and 3^′^10^′^, green). Significant separation over time is determined by a Kolmogorov-Smirnov (KS) test between the two profiles, identified to be at ≈ 28min. (e-g) Same as in (b-d), only for stochastic numerical simulations (n=50 repetitions) of the EGFR network shown in (a). (h) Top: Kymograph showing the differences in the network activity (Fig. 2a) over time for a reference trajectory for differentiation (3^′^20^′^) versus proliferation (3^′^3^′^) (mean trajectories from n=50 runs, respectively). Red and purple coloring indicates higher node activity in the differentiation and proliferation trajectory, respectively. Middle, bottom: averaged node activity from indicated periods (top panel) mapped to the network structure. The nodes Erk1/2, Bad, and Rab5 (highligthed in green) show the strongest differential response.

A challenge for verifying this concept experimentally is, however, that it relies on estimating the signaling trajectories that represent the overall activity of the higher-dimensional EGFR network over time. While this data is straightforward to obtain by mathematical modeling of the EGFR network, experimental measurements of the activity dynamics of a large number of proteins in the network in live cells with high temporal resolution are generally not feasible. In the aforementioned experiments, under each stimulus condition, the activity dynamics of a single MAPK protein, ERK, was measured with the biosensor EKAR2G in living cells over time^11^. To overcome this problem, we took advantage of a well-established method based on Taken’s delay embedding theorem^44^ from dynamical systems theory that enables the reconstruction of the overall activity of the signaling network from the information that the temporal dynamics of a single node carries about the rest of the network. Formally, the time delay embedding of a single time-series provides a one-to-one image of the original set on which the dynamics of the overall system evolves (cf. Methods, Supp. Figure 2a). Applying this approach, we obtained the signaling trajectories of the EGFR signaling network for the four distinct EGF signal frequencies from single cells (Figure 2b).

The signaling trajectories resulting from 3^′^3^′^ and 3^′^60^′^ EGF stimulation, both of which lead to proliferation in the experiments, display a strikingly similar, closed loop-like shape, despite high cell-to-cell variability. The trajectories induced by 3^′^10^′^ and 3^′^20^′^ stimulation leading to differentiation, in contrast, are characterized by repeated turns, but again, with a striking similarity between the two stimulation frequencies. To obtain a quantitative picture focused on the geometric shape of these trajectories, we employed dynamic time-warping^45^ before calculating the Euclidean distances between the obtained signaling trajectories. The 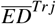 between signaling trajectories for proliferation-inducing (3^′^3^′^ and 3^′^60^′^) and differentiation-inducing EGF stimulus frequencies (3^′^10^′^ and 3^′^20^′^) remained low, confirming equivalent shapes between the trajectories within each class. In contrast, the 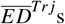 differed and significantly increased when comparing the signaling trajectories across the two classes (e.g. 3^′^3^′^ and 3^′^10^′^; 3^′^20^′^ and 3^′^60^′^, etc.; Figure 2c). These results confirm that signaling trajectories encoding for different phenotypes diverge in signaling state-space, enabling signaling specificity in a single network to arise even when stimulated with a single growth factor. Calculating, on the other hand, the 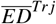 across time between the reconstructed space vectors of the signaling trajectories within and between classes showed that they significantly diverge at ≈ 28min from the start of the EGF pulse train (Figure 2d), suggesting this to be the earliest time point at which discrimination between the two distinct phenotypic responses can likely be detected from the signaling state of the network. Importantly, this implies that cell fate decisions under dynamic inputs occur rather rapidly after initial growth factor sensing: long enough for the cells to gauge the temporal signatures of the growth factors in their environment, but well before the signaling network sets to a steady state.

To obtain a deeper understanding of the underlying encoding mechanism, we next modeled the experimentally identified large-scale EGFR network (*N* = 56 proteins, Figure 2a)^42,43^ utilizing the biochemical system’s theory approach given by Eq. 2. We also accounted for the core features of the EGFR network that determine the temporal responses: a temporal memory in EGFR phosphorylation experimentally identified to be ≈ 40min in single cells^43^; as well as the effect of the EGFR spatial organization and related degradation, that re-sets EGFR phosphorylation to basal levels approximately one hour after single-pulse stimulation^14,43,46,47^. In the simulations, the network was stimulated equivalently as in the experiments, using temporal signals with 3^′^3^′^, 3^′^10^′^, 3^′^20^′^, and 3^′^60^′^ while mapping the resulting signaling trajectories. To mimic the experimentally observed cell-to-cell variability, we implemented stochastic simulations, with noise intensity in protein phosphorylation estimated from experimental measurements of temporal EGFR and ERK activity profiles by quantifying the correlation entropy^48^ from single-cell profiles (cf. Methods, Supp. Figure 2b). Plotting the average signaling trajectory obtained upon stimulation with the 4 distinct stimulus frequencies in a 3D representation (*EGFR*_*p*_, *MEKK*1, *ERK* chosen as examples) indicates overall similarities in their shape with the experimentally reconstructed signaling trajectories (compare Supp. Figure 3a with Figure 2b), although in the simulations, the kinetic rates were selected from uniform distributions, as the majority of these rates have not been experimentally determined in living cells. For comparison, we also used an experimentally parametrized *N* = 8 component EGFR-MAPK model^11,43^ and equivalently stimulated it with the four distinct signal frequencies. In this case, the obtained signaling trajectories showed a striking similarity to the experimentally derived ones (Figure 2e, compare to 2b). This shows that model parametrization is, of course, important for one-to-one mapping of the dynamics of the simulated to that of the physiological EGFR network, however, the *N* = 56 node network still captures the core features that contribute to the signal-to-phenotype mapping. Moreover, considering that the network dimensionality plays an important role in the separation of signaling trajectories in state space, as we have shown above (compare Figure 1c and Supp. Figure 1), we next performed the overall analysis on the *N* = 56 node EGFR network. The estimated 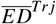 among signaling trajectories generated by the numerical simulations showed a significant difference only between the trajectories corresponding to proliferation and differentiation (Figure 2f), and the signaling trajectories encoding for different phenotypes diverged from each other ≈ 27min after the start of EGF pulse train in the simulation (Figure 2g). The simulations of the response dynamics of the *N* = 56 node EGFR network thereby qualitatively reflect the experimental data.

We therefore used this model to explore what is the mechanisms underlying the classification of the EGF frequency signals into distinct families of signaling trajectories, or in other words, how is the mapping of 3^′^3^′^ and 3^′^60^′^ (3^′^10^′^ and 3^′^20^′^) to signaling trajectories encoding for proliferation (differentiation) determined. The temporal traces of the node’s dynamics show that in the case of 3^′^60^′^ stimulation, the receptor and thereby the overall signaling state of the network remains active for approximately 40min after the 3min pulse, after which the network dynamics reset to the basal level. Thus, the trajectory makes one full rotation in the signaling state space before the stimulation with the second pulse in the sequence, giving the characteristic shape of the signaling trajectory (Supplementary Video 2). In the case of the 3^′^3^′^ stimulation, the system receives a series of 3min pulses (≈ 10) within the first hour of the stimulation, which in turn substantially depletes the active receptors on the membrane via internalization and unidirectional trafficking of the ligand-bound receptors (Supplementary Figure 3b). As a consequence, the system is effectively pushed towards the monostable regime (Figure 1c), which results in shortening or complete loss of the temporal memory. The EGFR phosphorylation, and thereby the overall activity in the network, is reset to basal level within the first hour from the start of the EGF stimulation (Supplementary Video 1). Hence, an equivalent excursion in the signaling state space is achieved as that for 3^′^60^′^ stimulation, explaining the similarities of the signaling trajectories in these two stimulation scenarios. In contrast, when stimulated with 3^′^10^′^ and 3^′^20^′^, subsequent pulses in the sequence arrive when the system is in the memory state, but spaced out widely enough in time such that EGFR concentration at the membrane only negligibly changes in the first hour of stimulation (Supp. Figure 3b). The upcoming pulses push the trajectory back towards higher activity in the signaling state space, thereby making the characteristic loops observed in the trajectories (Figure 2e, compare to 2b). Due to these state-space excursions driven by the EGF pulses, the trajectory does not fully reset to basal levels within the first hour of stimulation, resulting in a completely different trajectory shape, and thereby overall signaling state of the network. Thus, how growth factor signals are classified by the signaling network based on their frequency and mapped to a distinct family of signaling trajectories depends on the characteristic timescales of both the temporal memory of the core receptor circuit and the receptor internalization/degradation processes that together determine the trajectory shapes.

We next performed a complementary analysis on the signaling network dynamics from a network perspective. Using the 3^′^3^′^ and 3^′^20^′^ EGF frequency stimulation as representative signals that induce proliferation and differentiation, respectively, we estimated the differential activity of each node in the EGFR network during the first hour of the stimulation (Figure 2h). The differential map shows that around 27min, which we identified before as the earliest time point at which discrimination between the two distinct phenotypic responses can be detected from the signaling trajectories (Figure 2d, g), a sudden switch in the overall dynamics of the network occurs: the majority of the nodes in the network exhibit higher activity under 3^′^3^′^ as compared to 3^′^20^′^ stimulation (Figure 2h top, blue triangle, and middle). As time progresses, the overall signaling dynamics of the network is distinct between the two stimulus frequencies, with a majority of the nodes exhibiting higher activity under 3^′^20^′^ stimulation (i.e., *t* ∼ 45min), reflecting the further separation of the two signaling trajectory families. Interestingly, however, there are also time points where several nodes are distinctly activated under specific EGF signal frequencies (Figure 2h top, orange star and bottom). This implies that stimulation with a dynamic growth factor signal effectively induces a ‘modularization’ in time of a non-modular, distributed network. In other words, due to its dynamic properties, the nodes of the EGFR network are distinctly activated over time, thereby enabling specificity to emerge even for one growth factor signal. Interestingly, this phenomenon of temporal modularization seems to be most pronounced for three nodes of the EGFR network, ERK, Rab5, and Bad (Figure 2h, highlighted in green). Besides the well-known role of ERK during proliferation and differentiation, Rab5 is one of the main regulators of EGFR internalization and trafficking^47,49,50^, whereas previously, the Bad protein has been mainly discussed in the literature in relation to apoptosis^51^. Overall, this analysis suggests that simultaneous measurement of the temporal dynamics of these three proteins will likely yield maximal information on the signaling dynamics of the EGFR network when predicting phenotypic responses. Additionally, arranging the EGFR network topology to reveal its community structure using the greedy modularity community algorithm (cf. Methods) shows that these three proteins belong to three distinct communities (Supp. Figure 3c). Calculating the 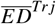 between signaling trajectories estimated from three nodes in the network showed that node selection from different communities aids the discrimination of signaling trajectories corresponding to different phenotypic responses in comparison to the selection of the three nodes from the same community, and this is significantly enhanced when using four proteins from four different communities (Supp. Figure 3d,e). This therefore suggests that experimental design should rely on measuring temporal protein activities from distinct communities to obtain predictive information on the cell fate decisions during the first hour of stimulation. Interestingly, we found that proteins belonging to the same signaling pathway, MAPK, belong to distinct communities. This is therefore in line with previously obtained experimental evidence that protein dynamics from a single pathway cannot be used to reliably predict the phenotypic response of the cells^10^.

Lastly, to confirm that homeorhetic regulation is a generic property of signaling networks irrespective of ligand/receptor identity, we performed the equivalent analysis for the experimental ERK activity time series for time-varying NGF stimulation (Supp. Figure 4). The trajectories induced by 3^′^3^′^, 3^′^10^′^, and 3^′^20^′^ NGF stimulus frequencies exhibited a similar multi-loop shape as previously observed for EGF stimulus frequencies inducing differentiation, whereas the trajectories obtained under 3^′^60^′^ NGF stimulation shared the single-loop shape with EGF-induced trajectories that lead to proliferation (compare Figure 2b and Supp. Figure 4a). The quantitative analysis of the similarity of the shape of the trajectories verified that the trajectories evoked by 3^′^3^′^, 3^′^10^′^, and 3^′^20^′^ NGF stimulus frequencies belonged to the same family, whereas 3^′^60^′^ NGF stimulation induced a different one (Supp. Figure 4b). Interestingly, the divergence in the signaling trajectories from the two families, and thereby the specificity in signaling, was achieved in this case much earlier, approximately 12min after initial NGF stimulation (Supp. Figure 4c). To model the NGF stimulation experiments, we accounted for the known differences in the spatial organization of TrkA receptors, which exhibit slower internalization in comparison to that of EGFR^47,52^, by decreasing their degradation rate. The simulations faithfully reproduced the experimental observations, including the time at which specificity in the overall signaling state of the network was achieved (Supp. Figure 4d-f). Furthermore, comparing the 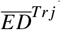 between the signaling trajectory families for proliferation and differentiation induced by both EGF and NGF (Supp. Figure 4g,h) showed that the growth factor signals encoding for a specific phenotype are mapped to a unique family of signaling trajectories, irrespective of the chemical identity of the signal, one of the main predictions of the novel concept os homeorhetic regulation of cell phenotype. Importantly, this supports the hypothesis that the existence of unique temporal activity patterns that can be generated by a single biochemical network under dynamic growth factor signals underlies distinct phenotypic responses.

Taken together the reconstruction of signaling trajectories from the experimental data on PC12 cells, the quantitative analysis of their shapes, and the complementary simulations demonstrate that the temporal signature of the growth factor, irrespective of the chemical identity, in conjunction with the memory and receptor characteristics of the signaling network generates and maintains a well-defined trajectory in signaling state-space that determines the PC12 phenotypic outputs in a homeorhetic manner.

### Robustness and flexibility of phenotypic responses is determined in *‘real-time’*

If the cellular phenotype is determined by the ‘flow’ of the trajectory in signaling state space, and if there is a unique mapping between the signal frequencies and the familly of signaling trajectories corresponding to a phenotypic response, the question naturally arises how robust the phenotypic responses are to changes in the stimulus frequency. Complementary to this is the question of how flexible the phenotypic responses are over time. Can cells adapt their response in real-time if the distribution of growth factor signals in the environment changes?

To gain a comprehensive picture, we used the model of the EGFR network (Figure 2a) to simulate the signaling dynamics over a set of EGF input frequencies between 3^′^3^′^ and 3^′^60^′^ and quantified the 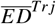 between the corresponding signaling trajectories and two reference signaling trajectories (3^′^3^′^ and 3^′^10^′^), thereby deriving which phenotypic response will be realized under each stimulus condition. The simulations show a switch for stimulus frequencies inducing proliferation to those inducing differentiation at 3^′^10, and in the opposite direction for frequencies above 3^′^40^′^ (Figure 3a, left panel). Interestingly, the trajectories corresponding to stimulation with 3^′^30^′^ and 3^′^40^′^ are equally distant to both reference trajectories for proliferation and differentiation, suggesting a possible third class of phenotypic responses that can be induced under EGF stimulation of PC-12 cells.

**Figure 3.**
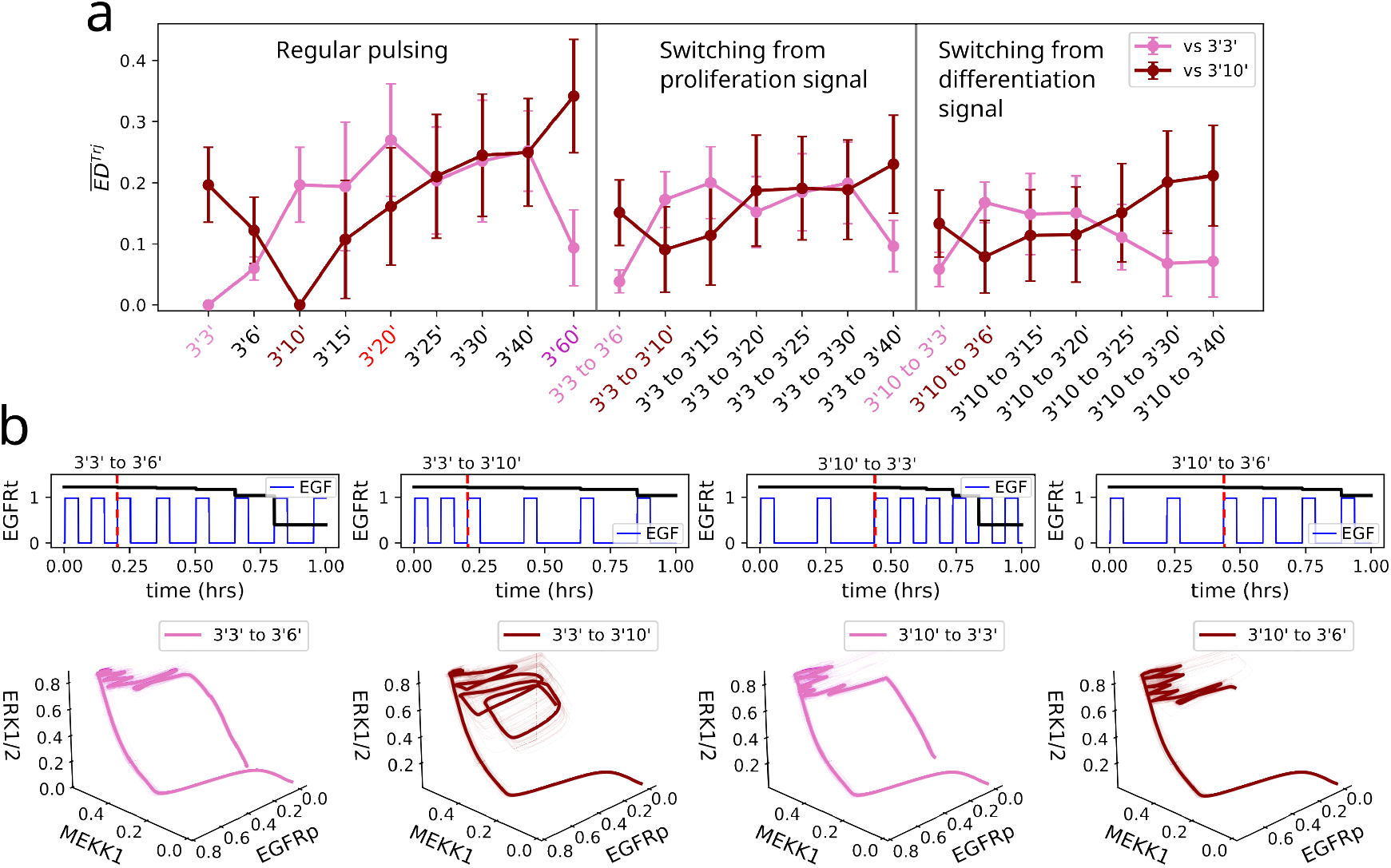
Real-time adaptation of phenotypic responses. (a) Euclidean distance (mean ±SD from N=40 stochastic simulation repetitions) estimated between the signaling trajectories generated by the EGF signal with composite frequency and by the reference signals for proliferation (pink) and differentiation (dark-red). (b) Exemplary representation of 4 distinct composite signals from the simulations, and the corresponding evolution of the total EGFR concentration (top), as well as the respective signaling trajectories (bottom).

To further test the robustness and hence the flexibility of the phenotypic responses when the temporal GF structure changes, we next stimulated the EGFR network with a complex EGF frequency profile: starting with two pulses corresponding to 3^′^3^′^ or 3^′^10^′^ EGF stimulation, the signal frequency was changed to another from the [3^′^3^′^, 3^′^60^′^] interval (Figure 3a, middle and right panel and examples in Figure 3b). In each condition, the 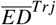 between the respective signaling trajectories and those generated by the reference signals was calculated for the first hour of stimulation. The quantification showed that robustness in the phenotypic responses is achieved not only for composite EGF signal frequencies belonging to the same class (i.e. the composite 3^′^3^′^*/*3^′^6^′^ frequency, as well as both frequencies independently induce proliferation, Figure 3b, left), but also across classes. For example, continuous stimulation with 3^′^6^′^ leads to proliferation, and with 3^′^10^′^ to differentiation; however, switching from 3^′^10^′^ to 3^′^6^′^ doesn’t alter the differentiation response (Figure 3b, right). The robustness of the cell phenotype for composite frequencies can be explained by the characteristic time-scales of the temporal memory in EGFR phosphorylation and of receptor internalization/degradation. Switching from a 3^′^10^′^ to a 3^′^6^′^ stimulation protocol does not effectively change the flow of the trajectory as the number of pulses and thereby induced amount of internalized receptors doesn’t significantly reduce the concentration of EGF receptor (Figure 3b, top right) and thereby the duration of EGFR phosphorylation memory, hence the initially encoded differentiation phenotypic response is preserved.

However, when stimulation with a composite frequency substantially changes the network characteristic in terms of available receptor concentration on the membrane, the phenotypic response can be dynamically adapted, as the cell processes the temporal changes of growth factor pulses from the environment. For example, switching from 3^′^3^′^ to 3^′^10^′^ will effectively induce differentiation (Figure 3b, middle left), whereas switching from 3^′^10^′^ to 3^′^3^′^ will lead to proliferation (Figure 3b, middle right). In the first case, switching from a 3^′^10^′^ to a 3^′^3^′^ stimulation frequency effectively changes the direction of the signaling trajectory in signaling state space as the increased frequency of pulsing, and thereby reduced concentration of internalized receptors (Figure 3b, middle left, top), pushes the system to reset to the basal state within the first hour upon stimulation beginning. This results in a closed broad-loop trajectory, similar to that for the reference proliferation signal. The opposite takes place when the stimulus frequency changes from 3^′^3^′^ to 3^′^10^′^. The decrease in the number of pulses throughout the first hour limits the internalization of receptors (Figure 3b, middle left, top). The transient memory duration is thereby preserved, enabling the signaling trajectory to dwell in the high activity areas of the signaling state space, resulting in the characteristic multi-loop signaling trajectory characteristic for differentiation. Lastly, the simulations predict that successful adaptation of the phenotypic response depends on a temporal “window of opportunity” in which the trajectory can alter its shape. For instance, changing the stimulation frequency after two cycles of 3^′^20^′^, or in total after 46min, to any EGF frequency from the broader set does not lead to a notable change of the trajectory within the first hour (Supp. Figure 5). This suggests that cells can potentially adapt their phenotypic responses to the signal changes in the environment only in a limited time frame after initial GF sensing. This time-frame depends on the initial frequency of stimulation and the characteristic time for trajectories separation for that growth factor (Figure 2d).

These results, therefore, demonstrate that homeorhetic regulation of signaling phenotype endows the systems both with robustness of the induced phenotypic responses, but also a necessary flexibility to adapt *on-the-fly*, when the temporal distribution of growth factor signals in the environment substantially changes. This is particularly important for cells that operate in dynamic environments, as it enables a real-time adaptation of their responses. From the perspective of cellular computations, this implies that biochemical networks in single cells are not only endowed with a signal classification property to uniquely map growth factor signals to families of signaling trajectories depending on their temporal characteristics, but can also dynamically regulate the responses in a homeorhetic fashion in non-stationary environments.

### Decoding information about phenotypic responses under dynamic signals resembles neuronal computations without stable states

Since both a commitment to a family of signaling trajectories, but also plasticity in the phenotypic response depending on the temporal signal distributions, are achieved within the initial hour of stimulus presentation, it is necessary to identify the dynamic requirements allowing IEG networks to decode this information in real-time. It has been experimentally demonstrated that promoters of IEGs are activated rapidly, generally with transient activation in normal cells^1,2,6^, giving the experimental basis for this hypothesis. Interestingly, it has also been shown that the same set of IEGs is activated by many different signals^55^, implying that the question of how specificity can be gained by activating the same components arises again, however, now on the level of gene regulatory networks. Considering this problem from a cellular computations perspective, this implies that the IEG network must possess the capability to distinguish and transform different internal states into given phenotypic outputs. In other words, the IEG network must continuously extract salient information from the high-dimensional trajectories generated by the signaling network and transform this information into a specific gene expression pattern.

An equivalent conceptual problem has been discussed on the level of neuronal networks: how a continuous stream of multi-modal inputs from a changing environment can be processed by stereotypical recurrent circuits of integrate-and-fire neurons in real-time. Utilizing the concept of reservoir computing (Figure 4a), it has been demonstrated that this task is reliably performed by a generic recurrent circuit having a fading memory represented by slow decay in the activity of the nodes of the so-called ‘liquid reservoir’, in conjunction with a memory-less feed-forward network that functions as a readout map, transforming at every time point the current liquid state into an output^53,54^. Making an analogy, the signaling network resembles the recurrent neuronal circuit as it is characterized by the presence of a temporal memory in the core receptor circuit. Hence, a feed-forward IEG network structure would likely be sufficient to enable decoding of the signaling trajectories to binary phenotypic outputs. Indeed, it has been identified experimentally that IEGs are induced in a conserved order over time, suggesting a feed-forward organization of the decoding layer^55^.

**Figure 4.**
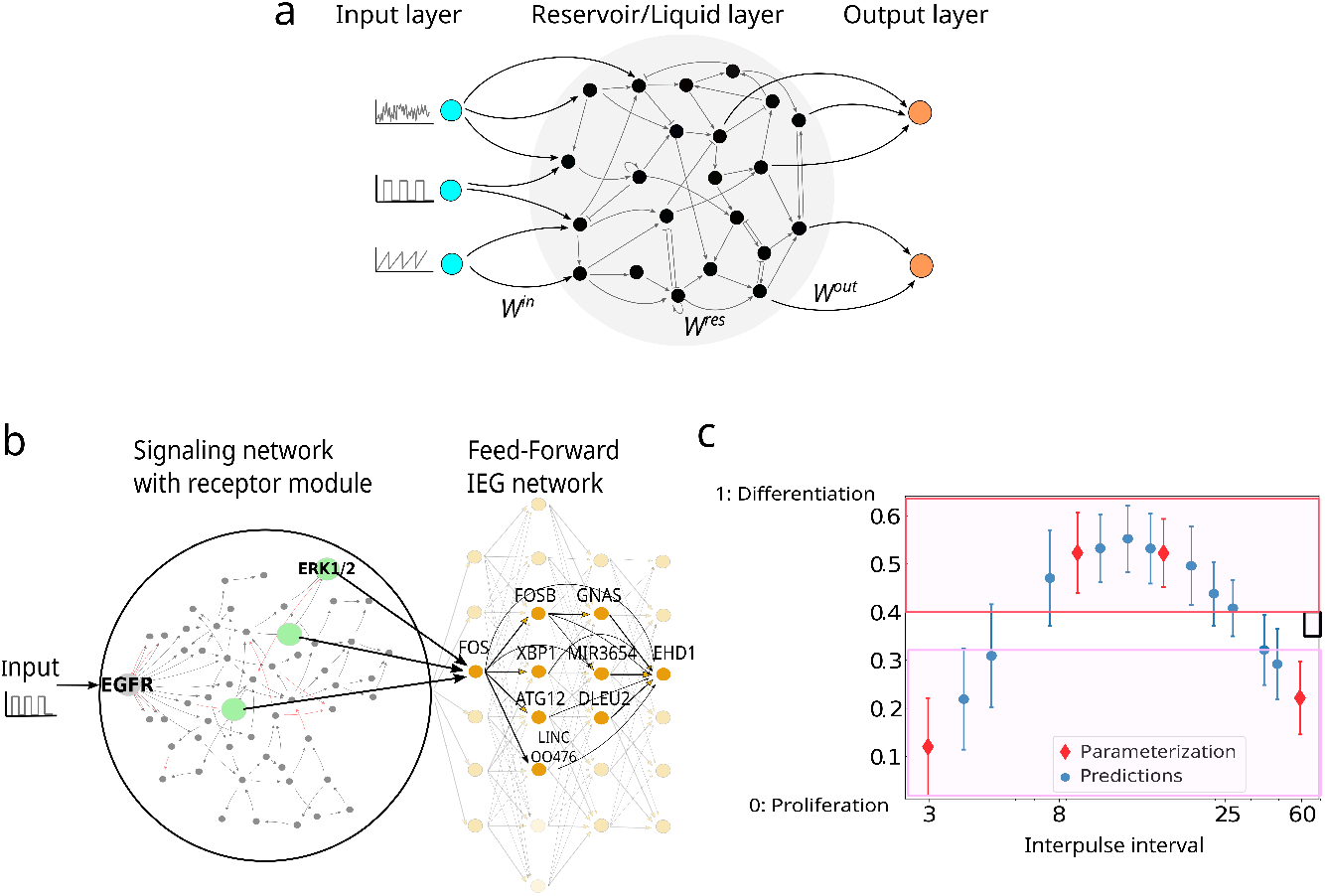
Decoding signaling trajectories with feed-forward immediate-early response gene regulatory networks. (a) Schematic representation of an artificial neural network (ANN), a neural liquid-state machine^53,54^, consisting of an input and output layers with feed-forward connectivity (*W*^*in*^,*W*^*out*^), and a recurrent reservoir (*W*^*res*^). (b) Schematic of the EGFR network and an experimentally identified IEG-network (orange nodes) arranged in a feed-forward structure. (c) Parametrization of the IEG network allows for salient information to be extracted from the high-dimensional signaling trajectory and transform it to stable binary readout, proliferation, and differentiation. Known signal frequencies are shown in red, and the predicted classification of unknown signal frequencies is shown in blue. Mean ±SD from N=10 stochastic simulation repetitions is shown.

We thus coupled the EGFR network (Figure 2a) to an ‘output’ feed-forward layer containing identified IEGs^55^: FOS, FOSB, XBP1, ATG12, LINC00476, DLEU2, MIR3654, GNAS and EHD1 (Figure 4b). Simple parametrization of the ‘output’ layer in terms of the weights of the connections, similarly as in the reservoir computing framework demonstrates that even this minimal IEG network architecture can effectively classify the signaling trajectories corresponding to the four reference signals (3^′^3^′^, 3^′^10^′^, 3^′^20^′^, 3^′^60^′^) into a binary output - proliferation (red points, magenta square in Figure 4c) or differentiation (red points, red square in Figure 4c), by inducing distinct temporal IEG activity patterns. Furthermore, this network can also classify a range of different signaling frequencies into one of the two phenotypes (blue points, Figure 4c), and the results are consistent with the predictions obtained from Figure 3a,b. Taken together, these results suggest that signaling homeorhesis enables robust encoding of cellular phenotypes on-the-fly, as the cell obtains information about the temporal distribution of the chemical signals in the environment, and feed-forward IEG network architecture is sufficient to decode this information in real time, thereby conceptually resembling how neuronal networks process time-varying signals.

## Discussion

Increasing experimental evidence shows that cells interpret in real-time the temporal distribution of growth factor signals in their multi-cellular environment to guide their phenotypic responses. In addition to the PC12 cells example^11^ discussed in this work, the frequencies of Wnt growth factor have been shown to directly influence cellular differentiation responses^29^, whereas frequency-dependent disruption of the Wnt and Notch dynamics, on the other hand, has been demonstrated to alter segmentation of the presomitic mesoderm during development^56^. In both cases, the presence of a slow time scale on the level of the dynamics of the underlying biochemical networks has been suggested as crucial. This slow dynamics likely arises from positioning in a critical vicinity to a dynamical transition, an infinite period bifurcation^57^. Such an organization is dynamically equivalent to the critical positioning close to bistability that results in temporal memory of the EGFR signaling network discussed here, only for oscillatory systems^58^.

This set of findings thus necessitates challenging the current view on how specificity in signaling networks is achieved. We have proposed here that a cell phenotype is encoded into a unique family of signaling trajectories, and distinct trajectories corresponding to different phenotypes diverge in the signaling state-space over time. The classification of signal frequencies to the trajectory families relies on the duration of the temporal memory of the core receptor circuit and the time scale of receptor internalization/degradation, in conjunction with the high dimensionality of the signaling network. This implies that context-dependent responses of cells are possibly generated by distinct temporal patterns of growth factors in different tissues, and that the different temporal characteristics of various receptors could be evolutionarily tuned to match the organ/tissue-specific frequencies of growth factor profiles. Signaling homeorhesis can thereby explain how the same growth factor can induce diversity of phenotypic responses, for example, proliferation, differentiation, apoptosis, etc., but also how chemically distinct growth factors can induce equivalent phenotypic responses while activating similar signaling network components. The framework also demonstrates that the IEGs likely decode the signaling state in real time, before the system settles to a steady state. One can, therefore, predict that the information about the phenotypic response can be obtained as early as ∼ 20min after initiation of the GF signal, as well as which protein activities in the network should be experimentally measured over time to reliably obtain this. The cellular computations underlying phenotype determination thereby resemble neuronal processing of a continuous streams of multi-modal inputs from a changing environment by stereotypical recurrent circuits of integrate-and-fire neurons in real-time^53,54^. The comparison to a neuronal network here is not only conceptual. In contrast, we find that it relies on similar principles: a recurrent signaling network that is endowed with temporal memory of the receptor circuit, as well as a feed-forward IEG network that is sufficient to decode the information from the signaling trajectories into binary outputs.

Moreover, the framework of homeorhetic regulation of cellular phenotype additionally allows for a description of how a balance between robustness and plasticity of phenotypic responses is established in dynamic environments. We show that cellular phenotype is determined in real-time, as the cell actively interprets the temporal signature of the growth factors in the environment, and the temporal characteristics of the receptor network, in particular the duration of temporal memory and the time-scale of receptor internalization and degradation, enable the coordination of this balance. Even more, we identified EGF signal frequencies that likely drive a third phenotypic response in PC-12 cells, a model prediction that needs to be further verified experimentally. Our conceptual framework thereby contrasts the current understanding that biochemical computations rely on a pathway-specific sensing of different ligand features, such as their molecular identity and concentration, which are transformed into specific concentrations and thereby steady-state responses of the intracellular effector proteins. Adding the layer of real-time cellular computations and encoding information through signaling trajectories rather than steady-states enables us to additionally explain how cellular phenotypic responses are determined under dynamic signals, but also how the phenotypic responses can be regulated in a homorhetic manner when the distribution of signals changes in the environment.

The term homeorhesis refers to “directional and distinct flow of the system’s dynamics or a system’s trajectory”, and has been initially introduced by the pioneer of theoretical biology, Conrad Waddington^39^, to describe canalization during development. He argued that under external changing signals, developmental systems tend towards an equilibrium; however, not one centered around a steady state, but rather on a direction or pathway of change, a description which is appropriate when the system doesn’t return to the same initial state. To bring a formal argument to this description, under dynamic signals, steady states are not the formal solutions of the system, as their number and position change; hence, the notion of steady states must be substituted with trajectories when investigating how cells process time-varying signals. We suggested here that the principle of homeorhetic regulation is much broader than initially proposed by Waddington, and applies in general to signaling regulation when growth factor signals are not constant over time. The framework of homeorhetic regulation of cellular phenotypes could thus be applied to study diverse biological processes, including development, regeneration, and cancer.

## Acknowledgements (optional)

The initial part of this work was performed at the Department of Systemic Cell Biology at the MPI of Molecular Physiology in Dortmund and we dedicate this paper to the memory of Phillipe Bastiaens, whose wisdom and creativity inspired this work. The authors thank Peter Bieling and the CCL group members for insightful discussions during the development of this work, and Robert Lott for critical feedback on the manuscript. A.K. acknowledges funding by the Lise Meitner Excellence Programme of the Max Planck Society. D.K. was funded by an EMBO Fellowship (Grant No. ALTF 310-2021).

## Author contributions

AK conceptualized and supervised the study, MY and DK performed the numerical simulations, and MY analyzed the experimental data. AK, MY, and DK interpreted the results. AK wrote the manuscript with the help of MY and DK.

## Code Availability

All code to reproduce the findings and figures from this study will be available under https://github.com/maneesh51/Homeorhesis upon final publication.

## Methods

### 1 Non-autonomous systems description of signaling trajectories

A complete trajectory is a particular example of an invariant set in a non-autonomous equation. In particular, a time-varying family of sets {Ξ(*t*)} is invariant if Ψ(*t, t*_0_)Ξ(*t*_0_) = Ξ(*t*), for all *t, t*_0_ ∈ ℝ.

### 2 Modeling of an arbitrary signaling network and signal classes generation

For the simulations in Figure 1, a signaling network with 30 nodes was considered, comprising a core receptor motive consisting of a double negative feedback loop^38^, that signals to a 27-node network with an arbitrary topology. The receptor network motif mimics typical receptor tyrosine kinase motifs^43,59–62^, whose parametric control ensures that the overall signaling network displays three distinct dynamical regimes: permanent memory (bistable organization), transient memory (organization at the critical transition between bistable and monostable regimes), or no memory (monostable organization). The bifurcation parameter (Figure 1c and Supp. Figure 1) in this case is the receptor concentration. The receptor motif is modeled using the law of mass action, and ensuring autonomous, autocatalytic, and ligand-induced receptor activation, where the receptor interacts by a double-negative feedback with a protein tyrosine phosphatase, the main regulators of receptor phosphorylation states:

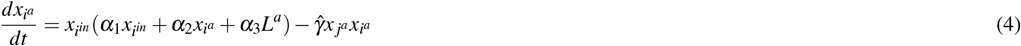

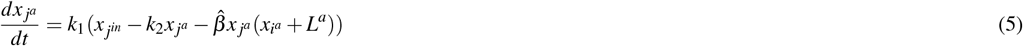

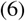

where 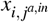 are the active and inactive forms of the receptor and the phosphatase, respectively, *L*^*a*^ represents the fraction of ligand-bound receptors. For binding/unbinding of ligand *L*_*T*_ to modulate *L*^*a*^ and additional term 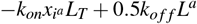 is added to the equation of 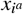, and the dynamics of *L*^*a*^ was modeled using 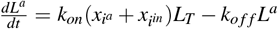. The 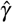 is the bifurcation parameter and is inversely proportional to the concentration of receptors on the membrane. The parameter values used are *α*_1_ = 0.0017; *α*_2_ = 0.3; *α*_3_ = 1; 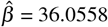, *k*_1_=0.01; *k*_2_= 0.5, *k*_*on*_=0.003 and *k*_*o f f*_ =0.01668.

The signaling network is generated using a probabilistic generation method, such that given *N* = 27 nodes, any two randomly selected nodes are connected with probability 0 *< p <* 1, assigning the sign of the link also with probability 0 *< p <* 1. The obtained network is modeled using biochemical systems theory (BST, Equation 2)^34,63^, where the activation and deactivation parameters are drawn from normal distributions.

The *reference* signal (*S*_*Re f*_) in Figure 1c (red profile) is generated with a fixed frequency, with a 3-minute pulse and 10-minute inter-pulse interval (3^′^10^′^). The classes {*S*_*I*_} and {*S*_*II*_} are generated in the following way: the position of each pulse in the sequence is shuffled with a probability *P*_*shu f f le*_ ∈ [0, 1], where *P*_*shu f f le*_ = 0 means all of the pulses are shuffled, and *P*_*shu f f le*_ = 1 - none. *S*_*I*_ class is generated with *P*_*shu f f le*_ = 0.95, and *S*_*II*_ with *P*_*shu f f le*_ = 0.1. Three exemplary input signals are shown in Figure 1c for {*S*_*I*_} (in blue) and {*S*_*II*_} (in green). Each class contains 10 input realizations. To estimate the similarity between the signals from *S*_*Re f*_ and {*S*_*I*;*II*_}, the Euclidean distance between each repetition from the *S*_*Re f*_ and {*S*_*I*;*II*_} was calculated as:

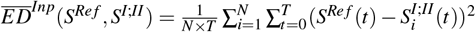

To estimate the contribution of the signaling network topology to the classification properties of the network (Figure 1d), the network described above is re-wired in the following way: given a linear cascade of nodes, a link between two nodes is removed/new link is added to a random node in the network with a probability 0 *< p <* 1, with an activatory (+1) or inhibitory (−1) sign also chosen with 0 *< p <* 1. We use these two network generation methods to obtain 15 different arbitrary network topologies of size *N* = 27, using the range of parameters given in Table 1.

**Table 1.** Network parameters for the cascade and rewire method (left) and the probabilistic method (right).

| Method 1 | Parameters |  |  |  | Method 2 | Parameters |  |
| --- | --- | --- | --- | --- | --- | --- | --- |
| | $P_b$ | $Prob(p)$ | $P_p$ | $P_n$ | | $Prob_p$ | $Prob_n$ |
| Net. 1 | 5/N | 0.5 | $1/N^2$ | $1/N^2$ | Net. 9 | 0.5/N | 0.5/N |
| Net. 2 | 15/N | 0.5 | $10/N^2$ | $1/N^2$ | Net. 10 | 0.1/N | 0.75/N |
| Net. 3 | 15/N | 0.5 | $10/N^2$ | $8/N^2$ | Net. 11 | 0.1/N | 0.5/N |
| Net. 4 | 15/N | 0.5 | $5/N^2$ | $2/N^2$ | Net. 12 | 0.1/N | 0.75/N |
| Net. 5 | 15/N | 0.5 | $5/N^2$ | $2/N^2$ | Net. 13 | 0.1/N | 0.75/N |
| Net. 6 | 18/N | 0.5 | $5/N^2$ | $2/N^2$ | Net. 14 | 0.75/N | 0.5/N |
| Net. 7 | 12/N | 0.4 | $5/N^2$ | $2/N^2$ | Net. 15 | 0.25/N | 0.15/N |
| Net. 8 | 12/N | 0.4 | $8/N^2$ | $2/N^2$ | | | |

For each of these networks, the following network properties are characterized: *N*_*A*_*/N*_*I*_ provides the ratio of activatory to inhibitory links, with 1 representing an equal number of links of both kinds, reciprocity provides a measure of the likelihood of nodes in a directed network to be mutually linked, and number of communities represents a subset of nodes within the network such that connections between the nodes are denser than connections with the rest of the network.

### 3 Modeling the 56-node EGFR network

The topology of the EGFR network shown in Figure 2a was obtained from^42^, and complemented with the EGFR-phosphatase interactions identified in^43^:

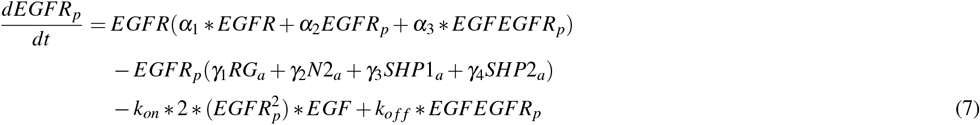

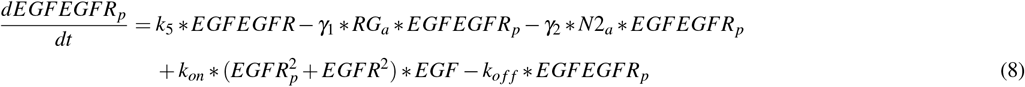

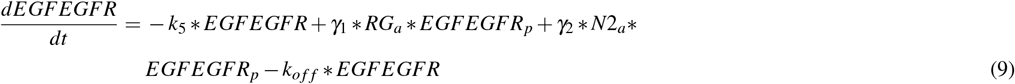

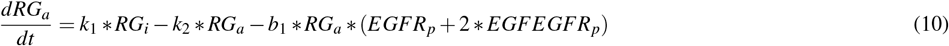

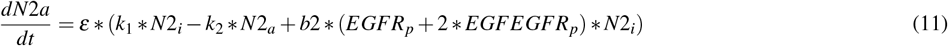

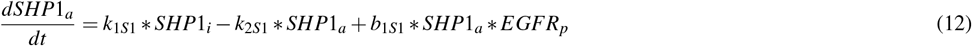

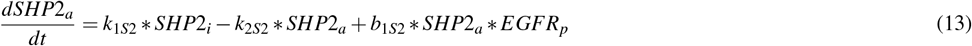

with *EGFR* = *EGFR*_*T*_ −(*EGFR*_*p*_ + *EGFEGFR* + *EGFEGFR*_*p*_), *k*_*d*_ = 5.56, *α*_1_ = 0.001, *α*_2_ = 0.3, *α*_3_ = 0.7, *γ*_1_ = 1.9, *γ*_2_ = 0.1, *ε* = 0.01, *k*_1_ = 0.5, *k*_2_ = 0.5, *k*_5_ = 1.613, *b*_1_ = 11, *b*_2_ = 1.1, *k*_*on*_ = 0.001, *k*_*o f f*_ = *k*_*d*_ * *k*_*on*_, *k*_1*S*1_ = 0.1, *k*_2*S*1_ = 0.9, *b*_1*S*1_ = 1, *k*_1*S*2_ = 0.15, *k*_2*S*2_ = 0.85, *b*_1*S*2_ = 1, *RG*_*i*_ = *RG*_*T*_ − *RG*_*a*_, *N*2_*i*_ = *N*2_*T*_ − *N*2_*a*_, *SHP*1_*i*_ = *SHP*1_*T*_ − *SHP*1_*a*_, *SHP*2_*i*_ = *SHP*2_*T*_ − *SHP*2_*a*_, *RG*_*T*_ = 1, *N*2_*T*_ = 1, *SHP*1_*T*_ = 1 and *SHP*2_*T*_ = 1.

The remaining EGFR signaling network was modeled utilizing Equation 2 from the main text, with parameters chosen randomly from uniform distributions, giving: 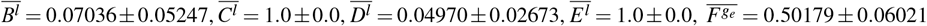, and 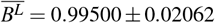.

### 4 Modeling the receptor-MAPK signaling module

To model the EGFR-MAPK network for Fig. 2e, the MAPK model from^11^ was coupled to the EGFR module identified in^38,43^, and also described and used in Figure 1. This corresponds to:

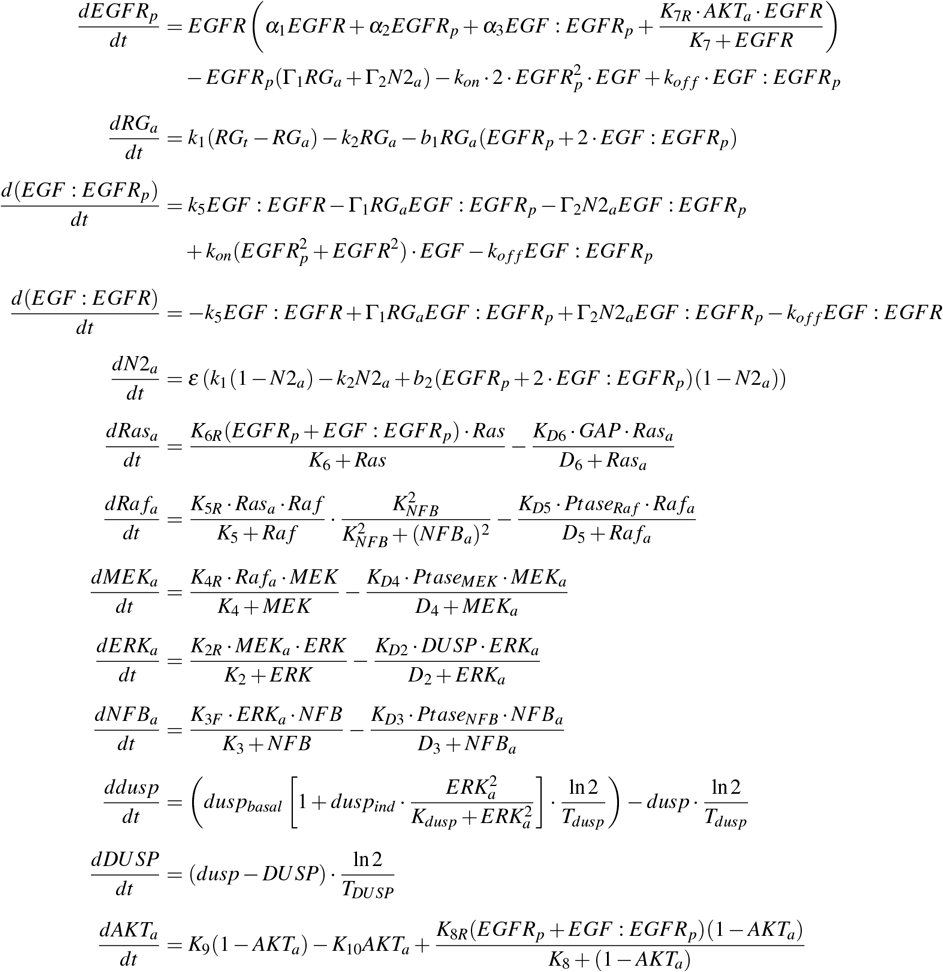

The model parameters are same as in^11,38^. To account for the degradation of total EGFR following signal pulses, as ligand-bound receptors are internalized and degraded^43,47^, a smoothed tanh-like sigmoidal decay function is used:

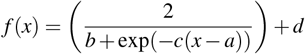

where *a* = 1.136, *b* = 2.53, *c* = 60, and *d* = 0.4. This form ensures a gradual but sharp decay of EGFRt after each pulse, moving the system away from the dynamical transition (Figure 1c), and hence reducing the duration of temporal memory. For stochastic simulations in all the figures, an additive noise term was added to each equation describing protein activity dynamics corresponding to a Gaussian white noise with zero mean and temporal correlation, and an noose amplitude corresponding to the experimentally identified ones.

### 5 Reconstructing phase-space trajectories from single-cell ERK activity recordings

We used the publicly available experimental data of temporal ERK response profiles obtained in single PC12 cells stimulated with EGF/NGF signals with diverse frequencies (for details of the experiments see^11^). The temporal ERK activity profiles from *N* = 30 cells per stimulation protocol were used. The stimulation protocols included stimulation with 3^′^3^′^, 3^′^10^′^, 3^′^20^′^ and 3^′^60^′^ 25*ng/ml* EGF and 50*ng/ml* NGF.

The reconstruction of the signaling trajectories was performed using the method of time delay, detailed as follows: For a time series of a scalar variable, a vector *x*(*t*_*i*_), *i* = 1, …*N* in state-space in time *t*_*i*_ can be constructed as *X* (*t*_*i*_) = [*x*(*t*_*i*_), *x*(*t*_*i*_ + *d*), .., *x*(*t*_*i*_ + (*m* − 1)*d*)], where *i* = 1 to *N* − (*m* − 1)*d*, and *d* is the embedding delay, *m* is a dimension of the reconstructed space, the embedding dimension. Following the embedding theorems by Takens^44^, if the sequence *X* (*t*_*i*_) consists of scalar measurements of the state of a dynamical system, then under certain genericity assumptions, the time delay embedding provides a one-to-one image of the original set, provided *m* is large enough. The minimum embedding dimension is obtained using *False Nearest Neighbor* method (FNN) proposed by Kennel et al.^64^, and the optimal time-delay (*τ*) is obtained using the mutual information (MI) as described by Fraser and Swinney^65^ using the `Python` package `skedm` (v0.1, https://skedm.readthedocs.io/en/latest/). The trajectory reconstruction was performed using a custom-made `Python` code. The single-cell ERK profiles were initially de-noised using `signal_savgol_filter` in `Scipy` module of `Python`. The average was taken over all cells per condition, and the MI and FNN were calculated, giving the respective time-delay and embedding dimension for each condition (Supp. Figure 3a). DTW was carried out using the *dtaidistance* Python package (https://pypi.org/project/dtaidistance/).

### 6 Estimating noise from time-series

To account for the noise levels characteristic for biochemical networks in our simulations, we estimated noise intensity from experimental measurements of temporal EGFR and ERK activity profiles. The noise estimation method is based on the functional dependency of coarse-grained correlation entropy *K*_2_(*ε*) on the threshold parameter *ε*^48^. The coarse-grained correlation entropy can be approximated using 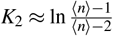, where ⟨*n*⟩ is the average diagonal line of length greater than 1. To estimate the noise level *σ* one can use correlation entropy *K*_*noisy*_(*ε*) and fit *K*_*noisy*_(*ε*)*ε* ^*p*^ to the corresponding experimental data. Therefore, one needs to estimate five free parameters *κ, σ, a, b*, and *c* for the function given by:

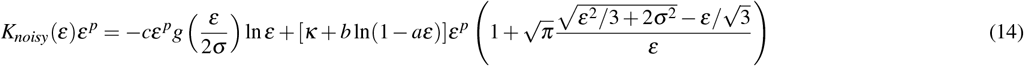

We applied this method first to *EGFR*_*p*_ temporal recordings obtained upon five minute EGF pulse stimulation of MCF7 cells (*N* = 30, exemplary profiles shown in Supp. Figure 2b; experimental methods as in^40^) to estimate the noise strength on the level of the core receptor circuit. First, each of the *EGFR*_*p*_ temporal profiles is used to obtain the entropy *K*_2_ using Eq. 11. Then, the correlation entropy *K*_*noisy*_(*ε*) curve is fitted to the experimental entropy using `curve_fit` function in `scipy.optimize` module of `Python`, as shown in Supp. Figure 2c. The parameter *σ* reflects the noise strength, here estimated to be *σ* = 0.023 ± 0.0133. Therefore, in the simulations, we use noise intensity 𝒪 (10^−2^) for the core receptor circuit. We also estimated the amount of noise present in the downstream signaling network by using the experimentally obtained *ERK* response profiles for PC12 cells (*N* = 30) stimulated with 3^′^3^′^ and 3^′^10^′^ EGF pulse trains (Supp. Figure 2d). Fitting the correlation entropy *K*_*noisy*_(*ε*) to the experimental entropy estimates (Supp. Figure 2e) yielded *σ* = 0.009 ± 0.0099, hence the noise intensity on the level of the signaling network that we used in the simulations was 𝒪 (10^−3^).

### 7 Parametrization of the IEG network

The IEG layer network is parameterized to generate an output of 1 for all phase-space trajectories corresponding to the signals inducing differentiation (i.e. 3^′^10^′^ and 3^′^20^′^ in case of EGF stimulation), and 0 for those inducing proliferation (i.e. 3^′^3^′^ and 3^′^60^′^). For this, we implemented the ‘reservoir computing’ framework^53^ from artificial neural networks, where the weights of the ‘reservoir’ in this case the protein-interaction network are not changed during training, and only the weighs in the output layer, in this case the IEG network, are trained to perform the mapping to binary output (1 differentiation, 0 proliferation). The training of the feed-forward networks was carried out using the Python library Tensorflow (https://www.tensorflow.org/about/bib).

## Supplementary information

### 8 Supplementary videos

**Supplementary video 1**. Evolution of the EGFR signaling network trajectory (purple) when stimulated with EGF 3^′^60^′^ (green dashed line, presence of EGF pulse) for the duration of the first hour after initial stimulus presentation. Corresponding to Figure 2e.

**Supplementary video 2**. Evolution of the EGFR signaling network trajectory (pink) when stimulated with EGF 3^′^3^′^ (green dashed line, presence of EGF pulse) for the duration of the first hour after initial stimulus presentation. Corresponding to Figure 2e.

**Supplementary video 3**. Evolution of the EGFR signaling network trajectory (pink) when stimulated with EGF 3^′^10^′^ (green dashed line, presence of EGF pulse) for the duration of the first hour after initial stimulus presentation. Corresponding to Figure 2e.

### 9 Supplementary figures and figure legends

**Supplementary Figure 1.**
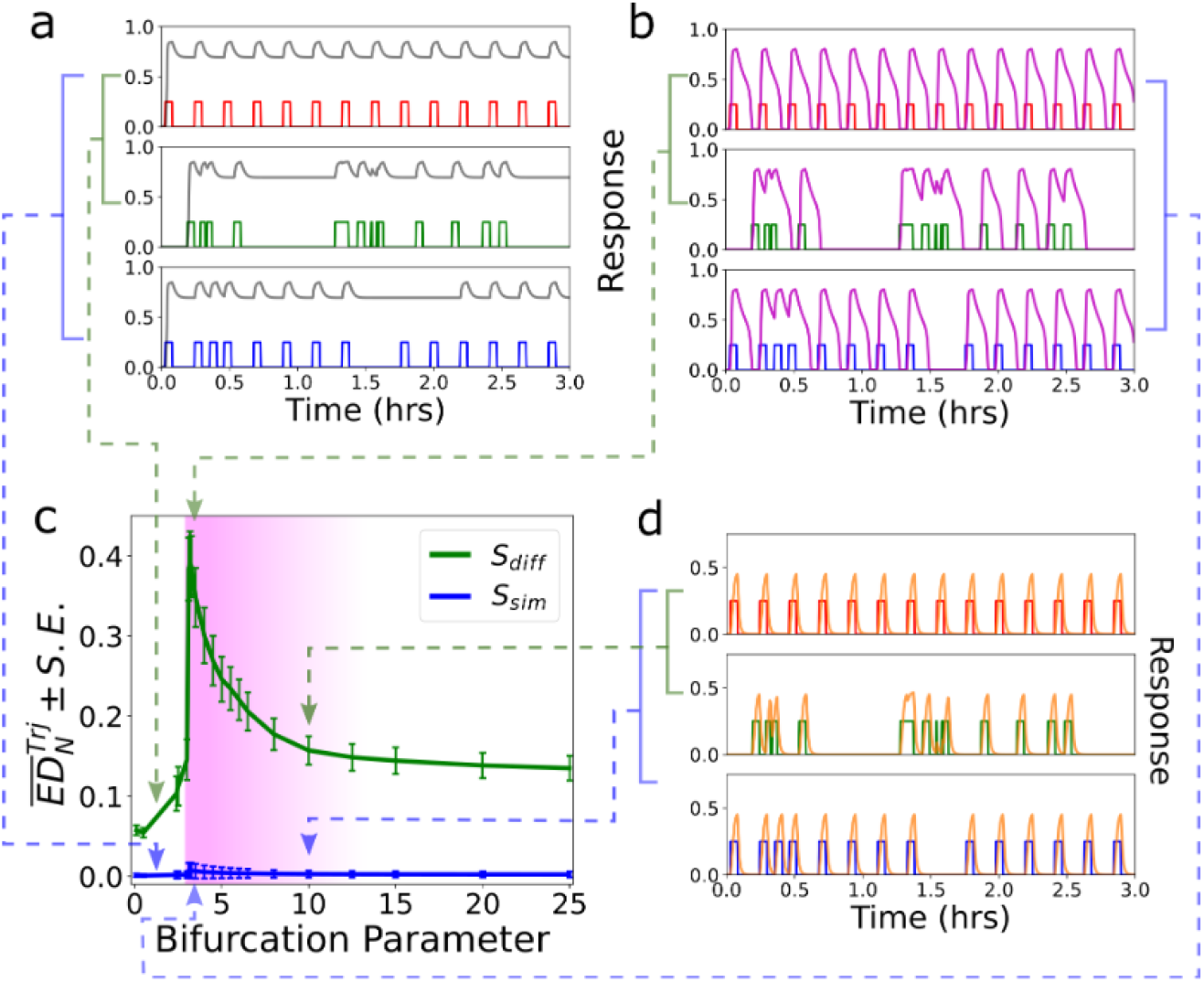
Signal classification with a minimal receptor module. Classification of signals based on their temporal profile by a generic receptor tyrosine kinase module (*N* = 3^38^) operating at three different dynamical regimes. Average 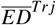 ± standard error between trajectories generated by 10 distinct inputs from *S*_*I*_ and *S*_*II*_ respectively, with respect to the *reference* trajectory in the 3-dimensional phase-space is shown as a function of the bifurcation parameter (receptor concentration). Insets show exemplary receptor phosphorylation profiles from the three distinct dynamical regimes for different signal frequency profiles.

**Supplementary Figure 2.**
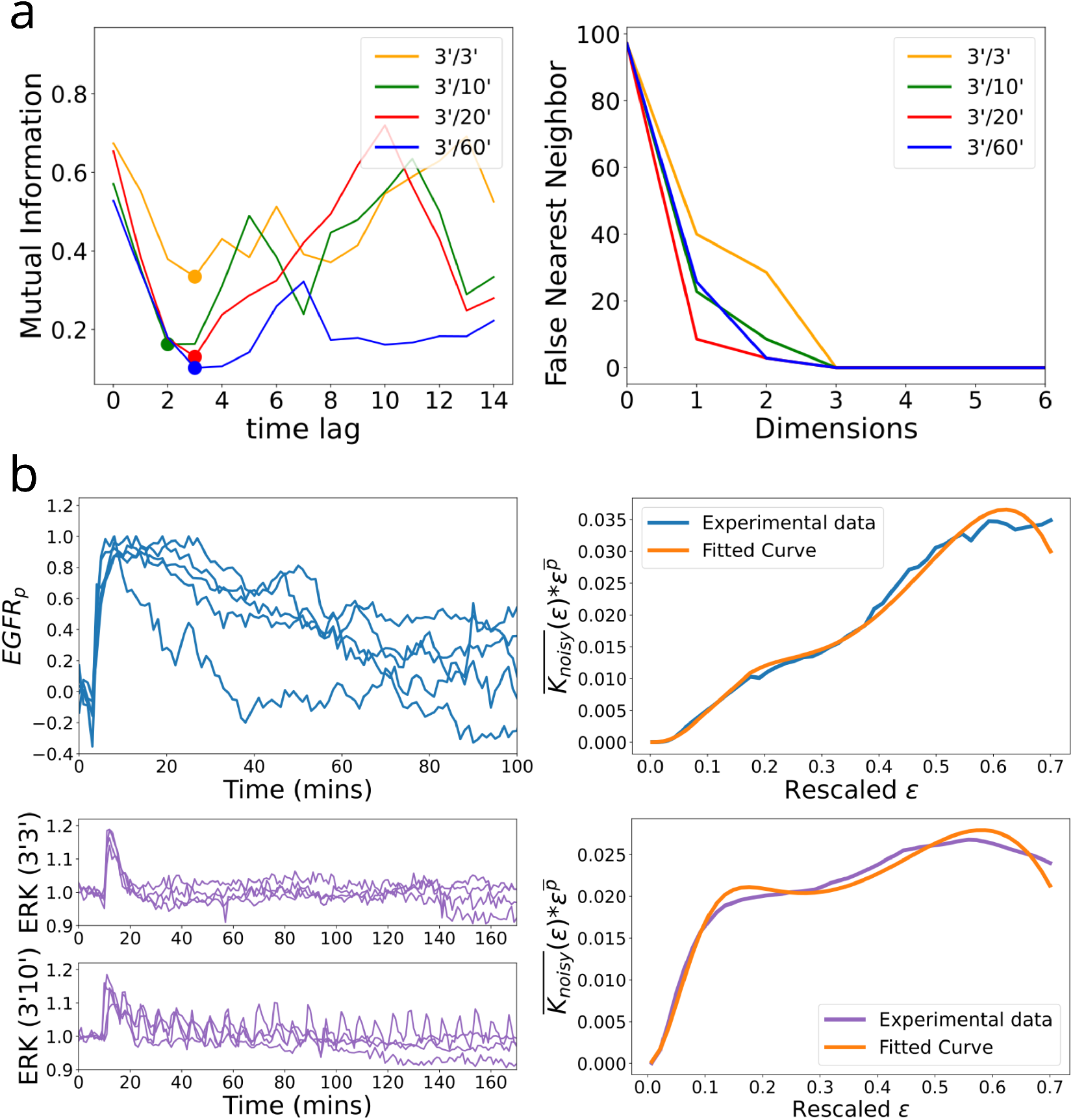
Reconstruction of phase-space trajectories and noise level estimation from experimentally identified protein phosphorylation time series. (a) Estimating the delay embedding using mutual information, calculated for the first one hour from experimentally measured temporal ERK profiles^11^ obtained under stimulation with 4 distinct growth factor frequencies. First *minimum* marked with dots of the same color as the respective signal frequency. Averages from *N* = 30 cells are shown. (b) Corresponding minimal embedding dimension obtained using *false nearest neighbor* method. (c) Exemplary *EGFR* phosphorylation time profile of 5 cells after stimulation by a single five-minute pulse of EGF stimulation. For experimental details see^40^ (d) *K*_*noisy*_(*ε*) function is fitted (in orange) for the entropy calculated using Eq.(11) for the *EGFR*_*p*_ time series profiles. (c) Exemplary *ERK* response time profile with EGF stimulation of signal frequencies 3’3’ (top) and 3’10’ (bottom). (d) *K*_*noisy*_(*ε*) function is fitted (in orange) for the entropy calculated from *ERK* data. The average *K*_*noisy*_(*ε*) value is taken over repetitions of cell numbers in both cases.

**Supplementary Figure 3.**
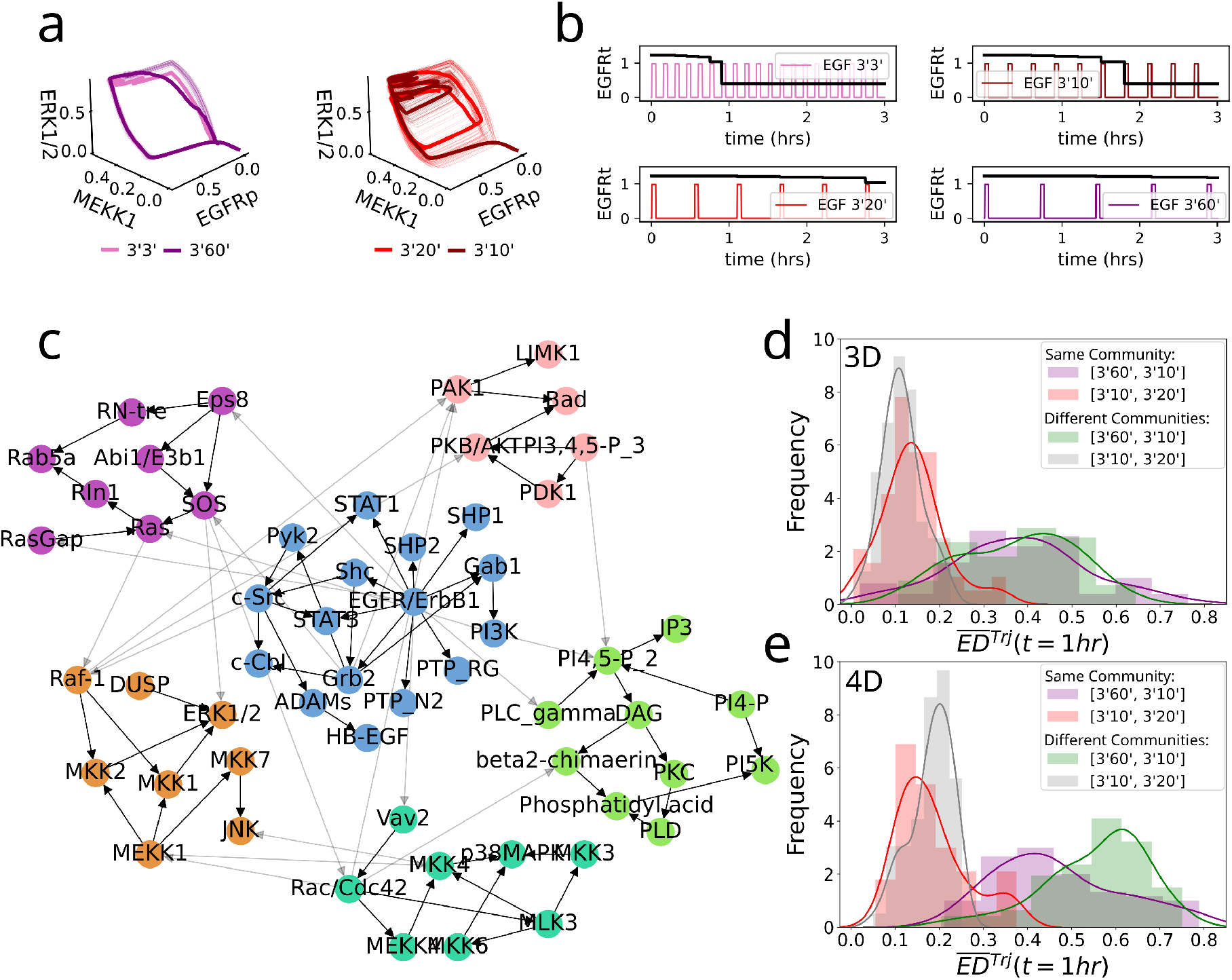
Characteristics of the EGFR signaling network. (a) Signaling trajectories obtained from stochastic numerical simulation of the EGFR network (*N* = 56, Figure 2a), depicting the overall evolution of the signaling network dynamics over time for different EGF input frequencies. Thin solid lines: single run of a stochastic simulation, thick solid lines: average per input frequency. Compare with Figure 2b and e. (b) Evolution of the total EGFR concentration over time under the different EGF stimulus frequencies obtained from the numerical simulations. (c) Community structure of EGFR network. 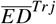 between trajectories obtained from a 3D (d) or 4D (e) representation, as estimated using temporal profiles from different nodes according to communities. 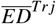 distribution when all the nodes are chosen from same (red and purple) vs. from different communities (grey and green) is shown.

**Supplementary Figure 4.**
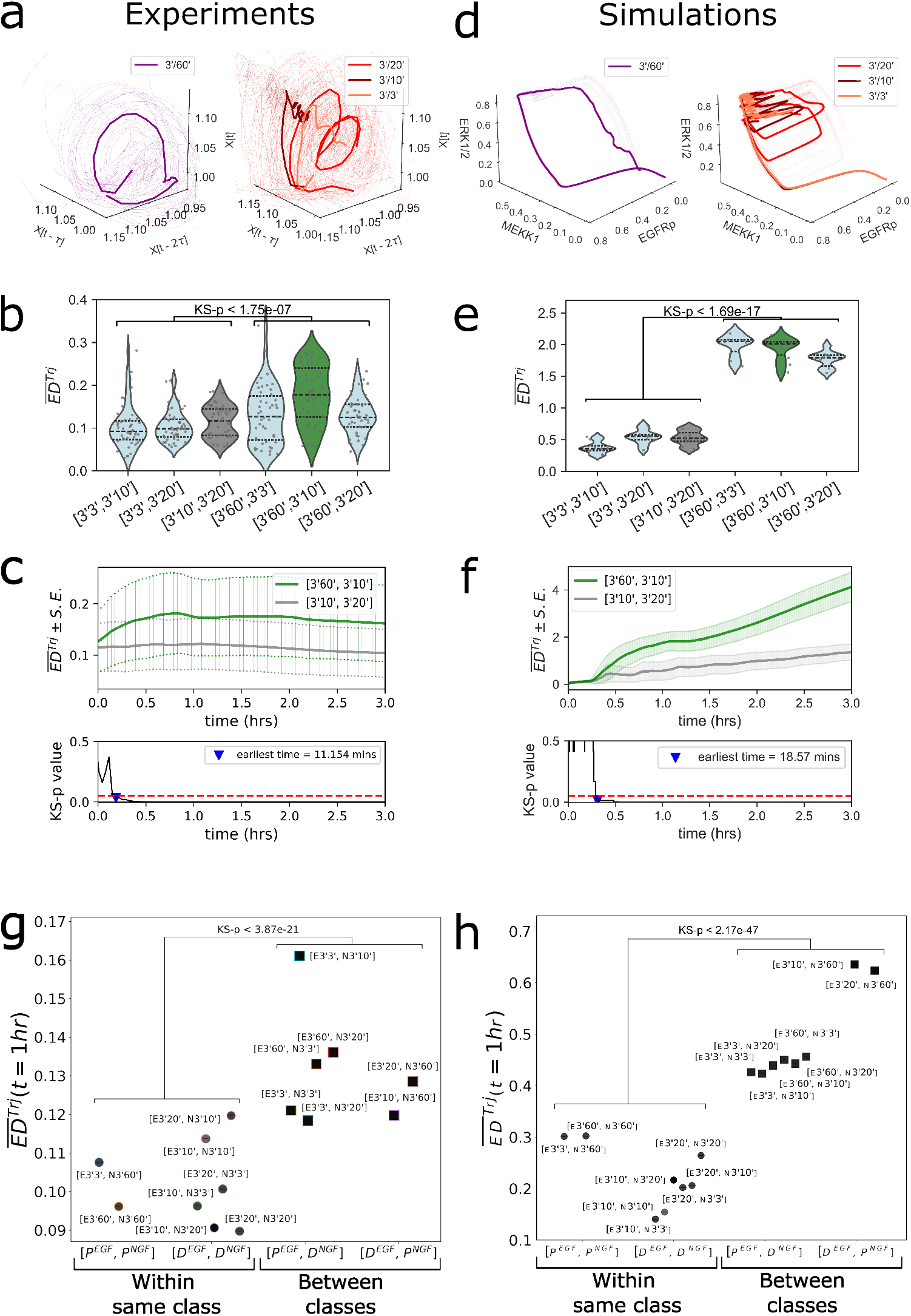
Homeorhetic regulation of cellular phenotype through distinct classes of signaling frequencies. (a) Reconstructed signaling trajectories from experimentally obtained temporal ERK phosphorylation profiles^11^, depicting the overall evolution of the signaling network dynamics over time for different NGF input frequencies. Thin solid lines: signaling trajectories reconstructed from single cell profiles, thick solid lines: average signaling trajectories per input frequency. (b) Mean Euclidean distance 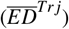 between signaling trajectories reconstructed from *N* = 30 cells per condition, calculated for the first hour of stimulus presentation, for all combinations of signals belonging to the same class i.e, proliferation and differentiation, and across the two classes. Trajectories were initially time-warped (Methods). (c) Evolution of the 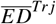 calculated between signaling trajectories reconstructed from *N* = 30 single cells corresponding to the same class (3^′^10^′^ and 3^′^20^′^, grey), or different class (3^′^60^′^ and 3^′^10^′^, green). Significant separation over time is determined by a Kolmogorov-Smirnov (KS) test between the two profiles, identified to be at ∼ 11min.(d-f) Same as in (a-c), only for numerical simulations of the TrkA receptor network (N=10 repetitions). (g) Mean 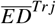 estimated from the reconstructed phase space trajectories (a-c and Figure 2b-d) for all trajectory combinations within and across classes. (h) Equivalent as in (g) fonly from numerical simulations data

**Supplementary Figure 5.**
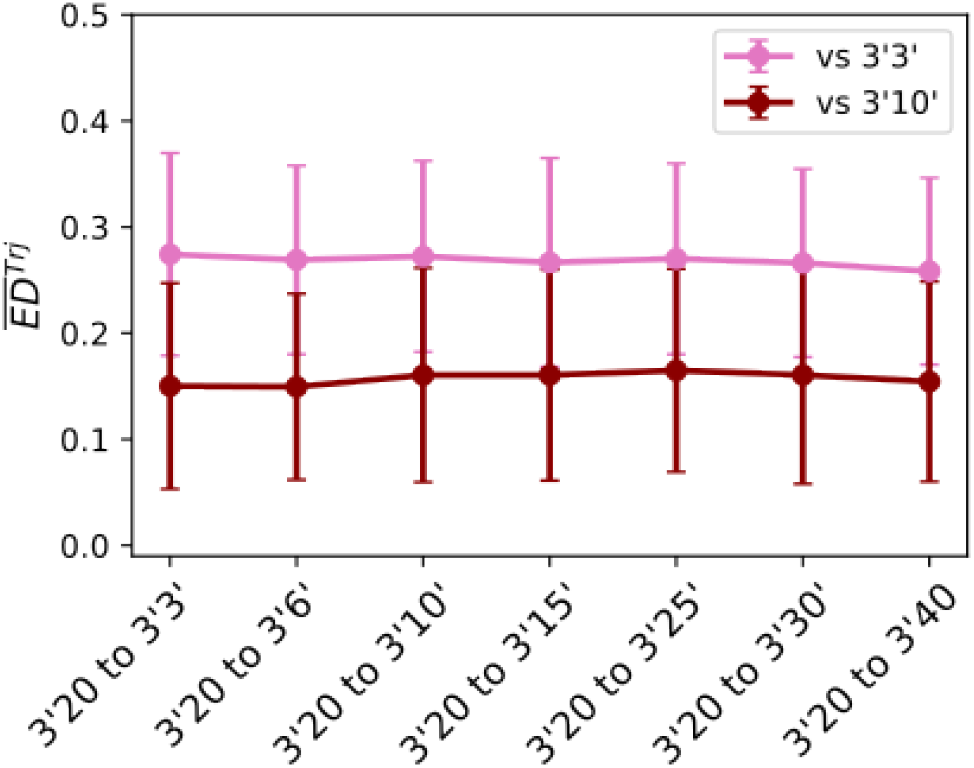
Robustness of phenotypic response to stimulation with composite frequencies. Euclidean distance (mean ±SD from N=40 stochastic simulation repetitions) estimated between the signaling trajectories generated by the EGF signal switching from 3^′^20^′^ to another EGF stimulation frequency. Note that two pulses of 3^′^20^′^ take 46 min, such that altering the shape of the trajectory within the first hour of stimulation is not possible even after a change of stimulus frequency. Thus, the 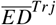 remains similar for all composite frequency profiles, leading to a robust phenotypic response.

